# Experimentally determined long intrinsically disordered protein regions are now abundant in the Protein Data Bank

**DOI:** 10.1101/2020.02.17.952028

**Authors:** Alexander Miguel Monzon, Marco Necci, Federica Quaglia, Ian Walsh, Damiano Piovesan, Giuseppe Zanotti, Silvio C. E. Tosatto

## Abstract

Intrinsically disordered protein regions are commonly defined from missing electron density in X-ray structures. Experimental evidence for long disorder regions (LDRs) of at least 30 residues was so far limited to a less than a thousand manually curated proteins. Here, we describe a comprehensive and large-scale analysis of experimental LDRs for 3,133 unique proteins, demonstrating an increasing coverage of intrinsic disorder in the Protein Data Bank (PDB) in the last decade. The results suggest that long missing residue regions are a good quality source to annotate intrinsically disordered regions and perform functional analysis in large data sets. The consensus approach used to define LDRs allows to evaluate context dependent disorder and provide a common definition at the protein level.

## Introduction

Despite a recent consensus regarding the existence of intrinsically disordered proteins (IDPs) and regions (IDRs) [1], its classification is still quite ambiguous [2]. As a result, various flavors of disorder have been proposed, some based on amino acid composition [3], flexibility [4,5] and functional roles coupled with conservation [6]. Perhaps the simplest distinction is between proteins with short and long disordered regions. Proteins with long disordered regions (LDRs) are special, since they seem to behave differently in function [2,7,8] and evolution [9].

Those regions which are poorly defined in the electron density map and consequently their position can not be defined are informed as missing residues. Missing residues in protein structures have been widely used as a proxy to identify IDPs/IDRs [10–12]. Nowadays, the PDB [13], the major repository of three-dimensional structures for proteins and nucleic acids, has more than 150,000 structures. PDB is mainly composed of X-ray (89%) and Nuclear Magnetic Resonance (NMR) (8%) structures, with a small number of Cryogenic electron microscopy (cryo-EM) (ca. 3%) and other techniques. A large scale analysis is possible by using high quality experimental data from thousands of protein structures. Structurally, disorder can range from regions that in solution are totally flexible to those that present two or more different, but defined, conformations [14]. Unfortunately, these two cases are often difficult to distinguish in an X-ray crystal structure, particularly at low or medium resolution. In fact, if a structure has a resolution better than 2.5 Å, it is possible to observe loops or short areas present in different conformations. However, at lower resolution these flexible regions are not visible in the electron density map. Consequently, the corresponding residues are left out from the molecular model. Cryo-EM also provides relatively high-resolution structures (with the exception of very few cases at a resolution better than 2 A, the large majority of them are at best 3 A) is being every year more abundant in PDB [15]. Contrary to X-ray diffraction, cryo-EM structures allow, at least partially, to distinguish the presence of LDRs from conformational flexible segments [16].

In this manuscript, those residues that are missing from the polypeptide chain (despite being present in the primary structure) are defined as “disordered”, without attempting to distinguish between disordered, flexible or mobile regions. Only long regions (at least 30 residues) were considered in order to disregard missing residues that may occur due to low resolution or experimental conditions. As different structures of the same protein may contain varying amounts of disorder, two consensus approaches were used to define unequivocally LDRs.

## Materials and Methods

### Long disorder data

UniProt [17] sequences with at least one structure in the Protein Data Bank (PDB) [13] (released until the 31st of December, 2018) were considered to perform this analysis. MobiDB [18] produced a list of 44,090 protein entries which could have structures coming from X-ray diffraction and/or cryo-EM. Cryo-EM structures were not considered in the present analysis.

Disordered residues at the UniProt protein level are those that are missing in the PDB structure. When more than one PDB structure is available for a given protein, two different consensus strategies were used, namely a “majority” and a “zero” rule. In the “majority rule” a segment is considered a LDR if it is disordered for at least 30 residues in the majority (more than 50%) of the structures corresponding to the same polypeptide chain. The “zero rule” is applied to the subset of majority cases where all the structures have a given LDR. Those protein fragments for which no PDB structures are available are considered as unknown/undefined.

Long disordered regions of at least 30 residues from DisProt (version 2019_08) [19] and IDEAL (release April 2019) [20] were used for comparison. The same majority rules were applied to IDEAL, and only disordered regions annotated with “disorder” and “high_rmsd” tags were considered.

### GO terms enrichment analysis

Functional enrichment was calculated for the first 4 levels of the Gene Ontology (GO) [21,22] graph as available in January 2020. Fisher’s exact statistical tests were carried out for the enrichment analysis using the LDR set as target (3,133 proteins) and all UniProt sequences with at least one PDB structure as background (40,200 proteins). A function was considered enriched if the p-value with Bonferroni correction was outside the 95% confidence interval of the mean (p < 0.05).

## Results and Discussion

One possible factor contributing to the presence of LDRs could be the resolution of the X-ray and cryo-EM experiments. In our dataset, we observe no correlation between the fraction of missing residues (disorder) per PDB chain and resolution. Indeed, Pearson’s correlation coefficients are 0.17 and −0.09 for X-ray and cryo-EM, respectively. It is important to mention that the way in which these techniques determine the resolution is different [23], and consequently their maps and models could not be equivalent [24]. In our data set, solely 2% of structures were obtained by cryo-EM. Hence, we only consider X-ray structures to perform the consensus disorder definition in order to avoid defining LDRs with missing residues coming from both techniques.

The release dates for the PDB structures in the dataset (see Figure 1b) show that most structures with LDRs have been deposited in the last five years, suggesting that the improvements of crystallization techniques is allowing to grow crystals of partially disordered or flexible proteins..

**Figure 1:**
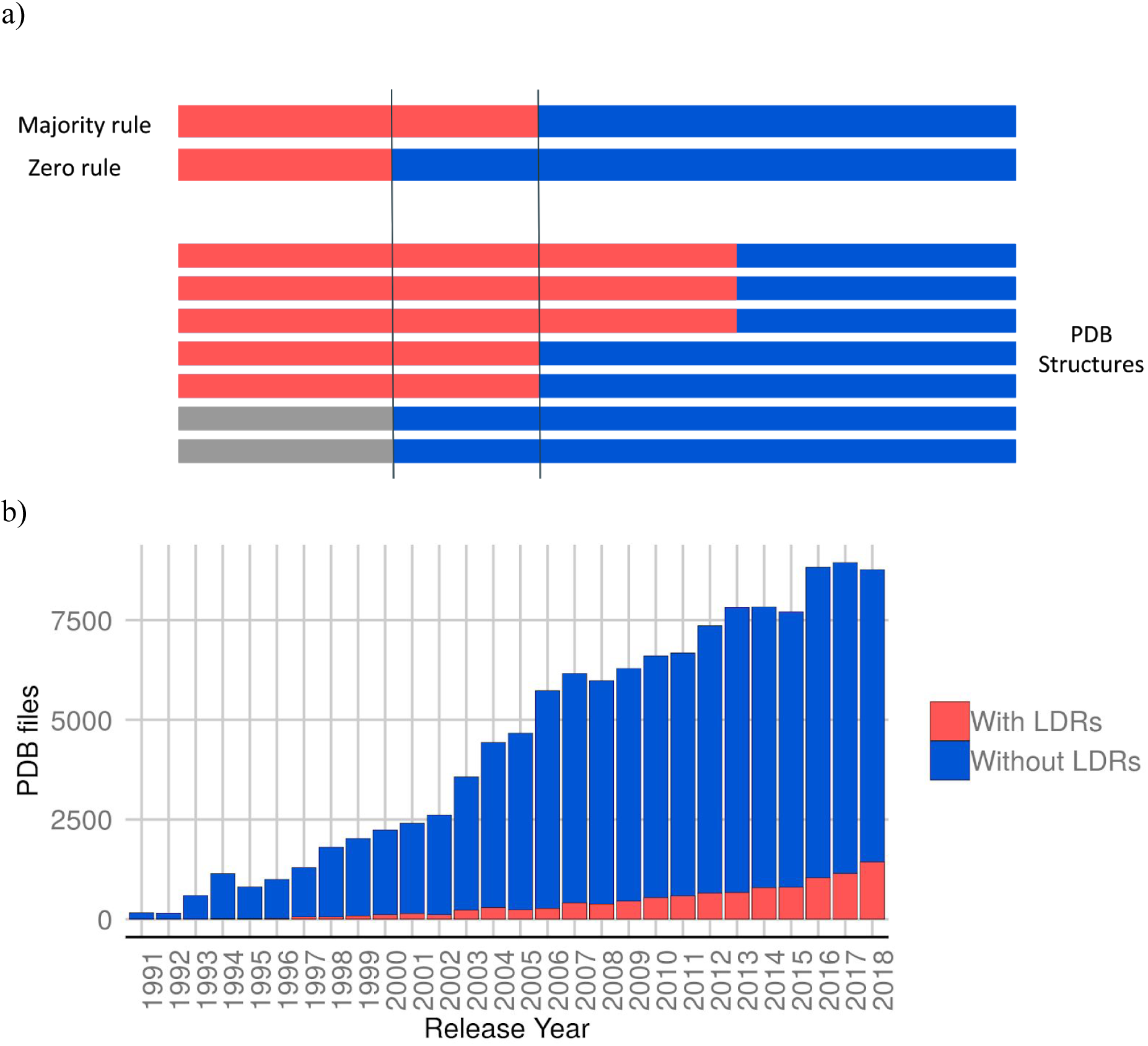
(a) Example of the majority and zero consensus definitions. A disorder consensus definition was applied when more than one structure was present for the same protein sequence. The color red represents the disordered residues (missing residues on the structure), blue refers to the structure part (ordered) and grey are those protein residues that do not have any PDB structure (unknown residues). Each bar on the top corresponds to a disorder consensus definition. Notice how the C-terminus is always disordered (zero consensus), while the first half has some X-ray structures suggesting order. Unknown residues do not affect positively or negatively to the consensus definition. (b) Amount of structures deposited by year in the PDB, highlighting the presence of LDRs (in red).

Figure 1a shows an example of the differences between the two consensus approaches, majority and zero rules. In total, the majority rule provided 3,133 proteins with at least one LDR, where 2,758 of them are confirmed with the stricter “zero rule” and 1,123 have a single PDB entry (See Table 1 for dataset composition). Other two sets could be derived from the initial dataset of proteins with at least one PDB structure. Those proteins without missing regions (completely structured proteins) and those proteins with disordered regions shorter that 30 residues (partially disordered). The disorder percentage of the LDR dataset (23.73%) is higher by design than other datasets [25]. The fraction of absent residues (26.5%) could harbour a further source of LDRs resisting crystallization, making our dataset a lower estimate for disorder.

**Table 1:** Dataset composition. The number of proteins, residues and LDRs is shown for the long disorder dataset, set of completely structured proteins (extracted for comparison) and partially disordered proteins (proteins with short disordered regions). Residues are unknown if there is no PDB structure assigned to those residues in the UniProt entry. More than one LDR per protein may be present.

| Dataset | Proteins | Residues |  |  | LDRs |
| --- | --- | --- | --- | --- | --- |
|  |  | Disordered | Structured | Unknown |  |
| Long disordered proteins | 3,133 | 270,656 | 1,140,513 | 509,174 | 3,553 |
| Completely structured | 8,366 | 0 | 1,829,962 | 1,130,817 | 0 |
| Partially disordered | 25,896 | 331,949 | 7,137,726 | 4,142,012 | 0 |

The distribution of LDR length is shown in Figure 2a, with DisProt [19] and IDEAL [20] for comparison, showing an exponential decay with increasing length. 50% of the regions of our dataset are between 30-44 amino acids and this decrease is consistent with IDEAL and DisProt. However, DisProt presents a bigger amount of extremely long regions (at least 200 residues) compared to IDEAL and LDR set. Our dataset has 93 proteins with these extreme LDRs which are a very unusual part of the PDB. Although each protein may contain more than one LDR, one region is the norm (2,773 proteins), with two being somewhat common (315 proteins). Combining the three sets will allow the construction of a larger set of 3,968 different proteins. Our data does not replace the excellent work of IDEAL and DisProt which are manually curated databases but rather offers a much larger and complementary experimental LDR source.

**Figure 2:**
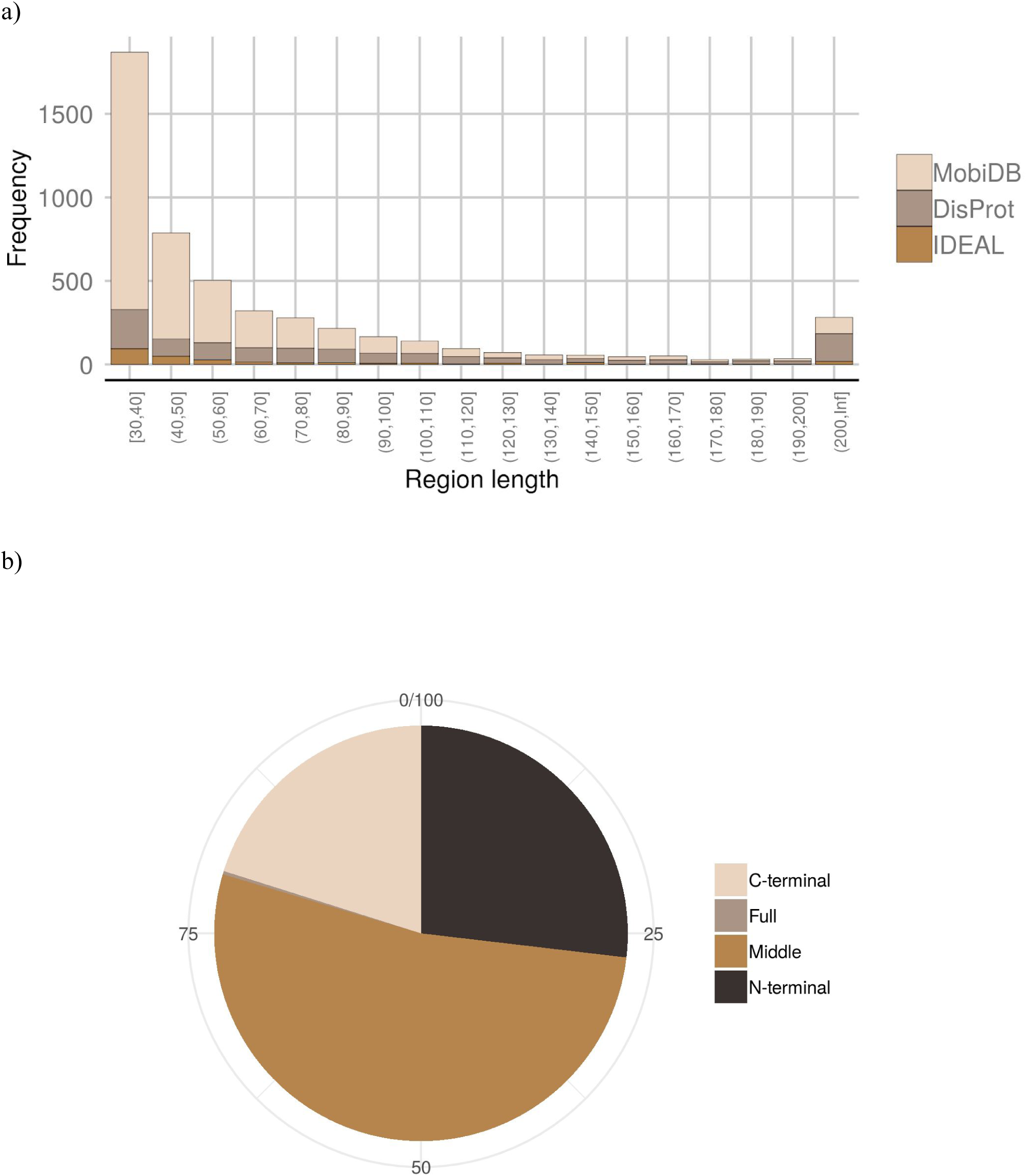
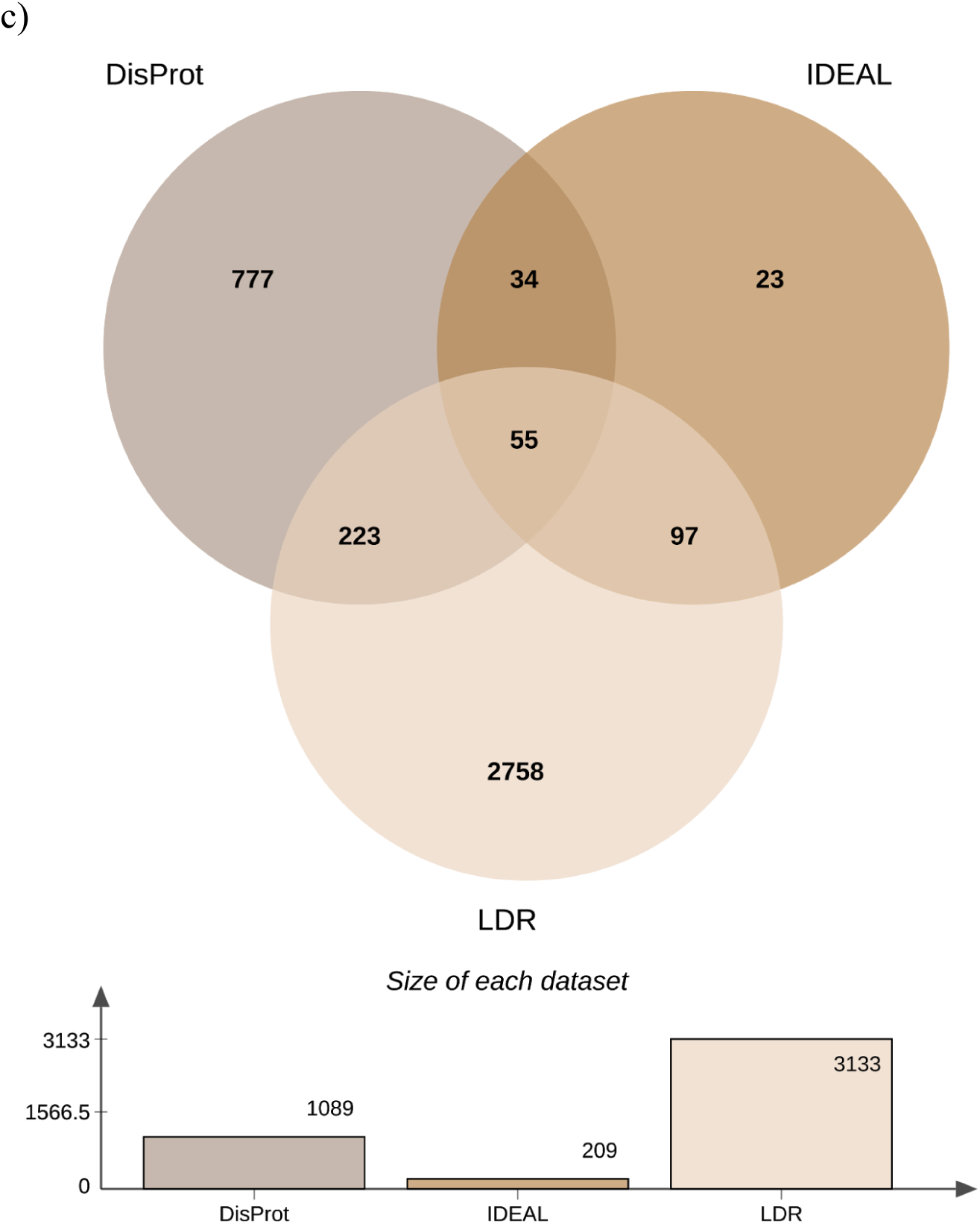
a) Length distribution of the disordered regions found in DisProt, IDEAL and LDR dataset. The data is grouped in bins of ten residues. (b) Fraction of LDRs which falls in N- and C-termini, full protein and the middle of the protein. Tails (N- and C-) refers to LDRs at the ends (20% of the total residues) of the PDB chain. (c) Venn diagram showing the overlap between the LDR dataset and the DisProt and IDEAL databases of manually curated disorder. Only proteins with LDR regions (at least 30 residues) were considered for IDEAL and DisProt databases.

While some of the LDRs may be the result of poor diffraction quality, it is well established now that the majority of them have functional roles [1,2,19,26,27]. To further support, we manually curated the 93 proteins with unusual LDRs by using the same curation procedure as we adopt for DisProt [19] (see Supplementary Table 1). 85% of LDRs have literature evidence that are disordered or unstructured, while 15% do not mention anything about the structural state of the region. Probably, missing residues were added during the structure refinement process for those proteins that we do not have any clue about disorder or simply the authors were not interested in characterise or mention the disordered region. Even some of the largest LDRs in X-ray structures are likely functional disordered regions instead of a result of specific or accidental experimental conditions, yielding a high quality dataset.

The size of the dataset allows us to perform function enrichments. We performed a Gene Ontology (GO) (Ashburner et al., 2000; The Gene Ontology Consortium, 2019) enrichment analysis to analyze the functional role of proteins with LDR, using as background all proteins with at least one PDB structure. The five most enriched terms in each ontology are shown in Figure 3. Intrinsically disordered regions (IDRs) function has been extensively studied in literature, not only in particular cases [2,28] but also in large scale studies [29,30]. Our LDR set is enriched molecular function terms, commonly associated with the IDPs/IDRs activity. The terms low-density lipoprotein particle binding and peptide hormone binding are related to the ability of IDPs to bind small molecules, macromolecules or other proteins. Protein prenyltransferase activity and protein deacetylase activity terms refer to the role of IDRs as effectors, interacting and modifying other proteins activities [2], while sigma factor activity is connected with the IDPs involved in transcription regulation. In biological process ontology, the LDR set is enriched in signal transduction terms (inositol lipid-mediated signal transduction and dopamine receptor signalling pathway), protein demethylation, biosynthesis and in the phosphatidylcholine metabolism. Among cellular components, protein with LDRs are mostly present in the nucleosome, chromosomes and protein-containing complexes. This last term is ancestor of three (cohesin, laminin and DNA packaging complexes) out five of the most enriched terms and could be associated with the capability of IDPs/IDRs to interact with different partners. In summary, many GO terms previously associated with disorder have been confirmed with our analysis and support the reliability of our LDR set [2,8,31,32].

**Figure 3:**
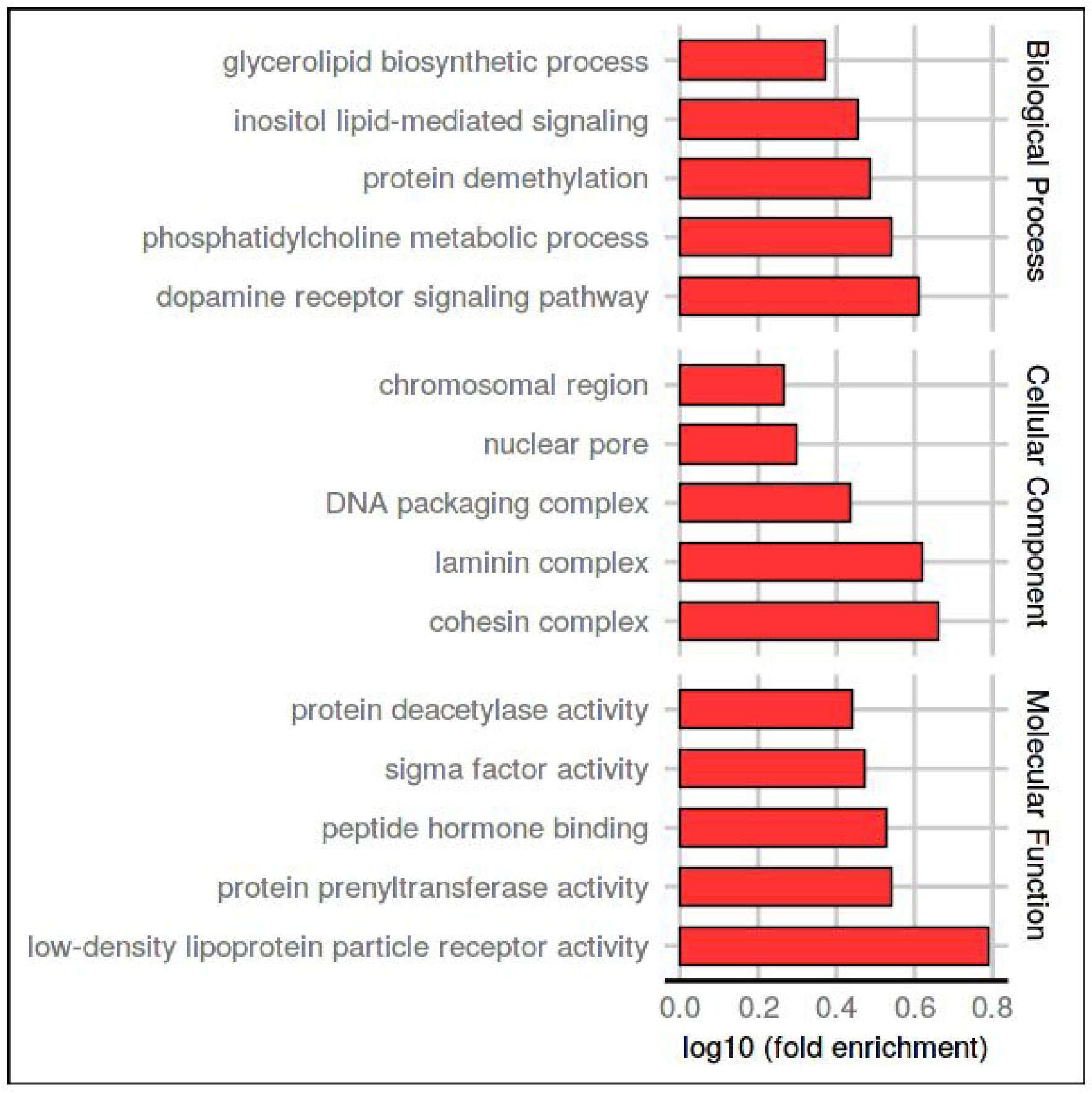
Five most enriched GO-terms in the three ontologies for LDR proteins. The background set was all UniProt sequences with at least one PDB structure. The x-axis shows the logarithmic increase compared to the background (see Methods for details).

We found 3,133 different LDR protein sequences from missing backbone atoms in X-ray structures. In case of relatively short regions, the missing electron density is often a consequence of alternative conformations in highly flexible areas, whilst for very long regions it most likely corresponds to unstructured portions of the polypeptide chain. The use of X-ray crystal structure in this study deserves a specific comment, since it is empirically well known that macromolecules with long flexible parts will tend to resist crystallization. It is common practice among crystallographers to produce different constructs of the same protein in order to reduce the flexible portions and, in doing so, favor crystal growth. In this sense, we would expect that our analysis underestimates the fraction of disordered regions present in the protein world. Most likely a larger fraction of disorder is present in the proteins that have not been crystallized yet. Most of the collective conclusions regarding long (and short) disordered regions have until now been based on predictions [25,30] and curated resources [7]. One of the main reasons for developing computational approaches was the scarcity of experimental data to make hypotheses. Despite this, predictors have given some interesting hypotheses with respect to LDRs, such as a functional analysis in full proteomes [8] and biological processes [33]. However, although predictors have good accuracy and can generate large quantities of data, they still contain systematic errors. For instance, on LDR proteins the predictor ESpritz [34] achieves 54.3% sensitivity and 91.2% specificity, while IUPred-long [35] produces a 38.9/93.6% sensitivity/specificity. While both prove a performance considerably above random, nevertheless substantial errors remain.

Our experimental LDR set is also significantly different from the currently available curated databases DisProt and IDEAL. It is important to stress that our data are not simply PDB entries, but rather multiple X-ray experiments assigned to frequently multi-domain UniProt sequences. Different X-ray experiments may be assigned to the same sequence with the final disorder/structure decision based either on majority evidence or complete lack of structure. This should produce a more stable definition since it will remove noise, e.g. missing residues arising from low resolution data or not well refined crystal structures. The majority definition also includes folding upon binding events (mixtures of disorder and structure), but we found these to be a rare occurrence as shown by the comparatively small difference (ca. 10%) between both definitions. However, more work is required to generalize the domains further, for example classifying them as “wobbly domains” or not. For a good example of a “wobbly domain” see Glutamine--tRNA ligase [36] from *D.radiodurans* (UniProt entry P56926) in our data set. It must also be considered that the presence of a domain whose orientation is flexible with respect to the rest of the protein may have two effects. One is to drastically reduce the probability of growing suitable crystals, the other is that the position of the wobbly domain becomes artificially frozen by the crystallization process, hampering its identification in the structure.

## Conclusions

A large dataset of diverse proteins with LDR is available to be used as a training set in disorder prediction techniques, as well as target IDPs to be included in the curated resources. Training a novel predictor on this large amount of quality data using state-of-the-art machine learning algorithms can only enhance our understanding of the phenomenon and improve their detection. Additionally, IDRs identified in this work could be used as a high-quality base ground to help in the annotation and identification of IDPs. A clearer picture will emerge as more structures are deposited each year in the PDB. Missing residues provide a valuable source of LDRs which tend to be overlooked in PDB as data source. In this work we demonstrate that PDB is not only the main repository of macromolecular structures but is also a good source to explore the (un)structure - function paradigm looking at disorder regions exposed to different experimental conditions, proving that most of the LDRs found have a biological role.

## Acknowledgments

This project has received funding from the European Union’s Horizon 2020 research and innovation programme under grant agreement No 778247 and No 823886. A.M.M. is funded by the research programme “MSCA Seal of Excellence @UniPD”.

## Abbreviations

IDR: intrinsically disordered regions;
IDP: intrinsically disordered protein;
LDR: long disordered region.

## Author Contributions

A.M.M., D.P. and M.N. collected the data sets; A.M.M. and M.N. performed the analysis; A.M.M. and F.Q. curated the data; A.M.M., M.N., I.W., D.P., G.Z. and S.T. designed the study and wrote the manuscript; S.T. conceived the idea.

